# gTranslate: rapid and accurate translation table prediction for prokaryotic genomes

**DOI:** 10.64898/2026.05.24.727570

**Authors:** Pierre-Alain Chaumeil, Philip Hugenholtz, Donovan H. Parks

## Abstract

**Background:** Bioinformatic tools often require the prediction of protein-coding genes to make inferences about prokaryotic genomes. Typically, the genetic code used for translating genes to proteins must be specified by the user based on the taxonomic classification of a genome assembly or, for some widely used tools, established using a heuristic rule based on gene coding densities. Manual specification is at best inconvenient, but more challenging is that many bioinformatic tools are applied before taxonomic classifications have been established making specifying the translation table impractical.

**Methods:** Here we provide a computationally efficient tool, gTranslate, that uses an ensemble of five machine learning methods to accurately predict translation tables for prokaryotic genomes. The feature vector used by gTranslate takes advantage of differences in gene coding densities when predicting genes under different translation tables along with features that consider the number and ratio of UGA stop codon reassignments to tryptophan or glycine.

**Results:** We demonstrate that gTranslate correctly predicts the translation table of prokaryotic genomes >99.99% of the time (i.e. <1 error per 10,000 genomes) and outperforms a more computationally expensive prediction method and a coding density heuristic used by popular bioinformatic tools. Using gTranslate, we identify a basal lineage of *Ca*. Stammera capleta that uses the standard bacterial genetic code instead of the UGA stop codon to tryptophan reassignment common to other members of this species. We also identify the first instances of UGA-to-tryptophan reassignment in the *Patescibacteriota* making this the first bacterial phylum with members capable of using translation tables 4, 11, and 25.

## Introduction

The genetic code defines the correspondence between nucleotide triplets (codons) in messenger RNA and the amino acids or stop signals they specify during protein synthesis (Koonin and Novozhilov, 2009). Although this code is nearly universal across all domains of life, deviations from the standard code have been documented in both nuclear and organellar genomes spanning eukaryotic, bacterial, and archaeal lineages (Knight et al., 2001; Shulgina and Eddy, 2021). These deviations most commonly involve the reassignment of stop codons to amino acids, though sense codon reassignments have also been identified in certain eukaryotic lineages (Mühlhausen and Kollmar, 2014) and have been computationally predicted to occur in some prokaryotes (Shulgina and Eddy, 2021). Despite the diversity of known reassignments, the high conservation of the genetic code across billions of years of evolution underscores the significant selective constraints on its modification and the rarity of successful reassignment events.

While more than 99% of known prokaryotic species use the standard bacterial genetic code i.e. translation table 11, two variant codes have been identified in bacteria involving reassignment of the UGA stop codon. Translation table 4 encodes UGA as tryptophan and is used by all species of the *Bacillota* order *Mycoplasmatales*, a group of endosymbionts primarily associated with vertebrates, arthropods, and plants (Razin et al., 1998). This reassignment also occurs in several *Pseudomonadota* endosymbionts with highly reduced genomes, including *Candidatus* Hodgkinia cicadicola in cicadas (McCutcheon et al., 2009), *Ca*. Zinderia insecticola in spittlebugs (McCutcheon and Moran, 2010), *Ca*. Nasuia deltocephalinicola in leafhoppers (Bennett and Moran, 2013), *Ca*. Stammera capleta in tortoise leaf beetles (Salem et al., 2017), and an *Fastidiosibacteraceae* species in dinoflagellates (Nakayama et al., 2025). More recently, this reassignment has been identified in two *Verrucomicrobiota* ciliate endosymbionts *Ca*. Organicella extenuate (Williams et al., 2021) and *Ca*. Pinguicoccus supinus (Serra et al., 2020), and in two genera in the *Actinomycetota* family *Eggerthellaceae* (Parks et al., 2025). Translation table 25 encodes UGA as glycine and was first identified in members of the candidate phylum *Absconditabacteria* (Campbell et al., 2013), with this recoding also found in *Gracilibacteria* (Rinke et al., 2013). These bacterial reassignments share several genomic hallmarks, including reduced genome size, compositional bias, and loss of the peptide chain release factor RF2 (McCutcheon et al., 2009). In addition, many lineages with a stop codon reassignment contain tRNA^Trp^ or tRNA^Gly^ with the anticodon UCA to directly facilitate translation of UGA to tryptophan or glycine, respectively (McCutcheon et al., 2009; Nakayama et al., 2025). The first genome-wide stop codon reassignment in archaea was recently reported; a UAG-to-pyrrolysine reassignment predicted in multiple archaeal lineages and validated by proteomic evidence (Kivenson et al., 2025).

The genetic code used by an organism can be established from its taxonomic classification, but this presents a practical challenge for bioinformatic workflows that require protein-coding gene predictions before taxonomic classifications have been established. A primary example is GTDB-Tk, which classifies genomes relative to the Genome Taxonomy Database (GTDB) by placing them into a reference tree based on the concatenation of phylogenetically informative protein sequences (Chaumeil et al., 2022). More broadly, prediction of protein-coding genes and their correct translation is an essential step in many common bioinformatic applications, including estimating genome quality (Chklovski et al., 2023; Manni et al., 2021), annotating the function of proteins (Seemann, 2014; Tanizawa et al., 2018), and taxonomic profiling of metagenomic samples (Dmitrijeva et al., 2025; Woodcroft et al., 2025). Tools have been developed to detect genetic code reassignments in phage genomes (Pfennig et al., 2023) or from eukaryotic sequence data such as GenDecoder that targets metazoan mitochondria (Abascal et al., 2006), Bagheera that predicts CUG codon translation in yeast species (Mühlhausen and Kollmar, 2014), and CoreTracker that detects codon reassignments in mitochondrial genomes using a phylogenetic framework (Noutahi et al., 2017). FACIL and its successor Codetta provide a more general approach by aligning nucleotide sequences against profile hidden Markov models of conserved protein domains to infer the most likely amino acid encoded by each codon (Dutilh et al., 2011; Shulgina and Eddy, 2021). However, while these tools provide evidence of potential codon reassignments, they do not make explicit predictions regarding which translation table a genome uses and are computationally expensive as they require homology searches against large protein databases.

Bioinformatic tools often require users to specify a translation table, which is at best inconvenient, but often impractical to determine resulting in prokaryotic genomes being erroneously processed with the standard bacterial genetic code. This results in artificially truncated and potentially misannotated proteins in lineages with a stop codon reassignment. More recent tools address this issue by using a coding density-based heuristic to predict the translation table of prokaryotic genomes such as CheckM (Parks et al., 2015). The CheckM heuristic considers the coding density of genomes using the standard prokaryotic genetic code (CD_11_) and UGA recoded for tryptophan (CD_4_), relying on the observation that genomes recoding UGA to an amino acid will have atypically low coding density when genes are predicted using translation table 11. Specifically, this heuristic uses the decision boundary CD_4_ – CD_11_ > 5% and CD_4_ > 70% to establish a genome as using table 4. This approach has been adopted by several widely used tools, including GTDB-Tk (Chaumeil et al., 2022), CheckM2 (Chklovski et al., 2023), and SingleM (Woodcroft et al., 2025). Similar coding density-based strategies are used by BUSCO (Manni et al., 2021) and MiGA (Rodriguez-R et al., 2018).

While coding density-based heuristics generally perform well, even a low misclassification rate becomes problematic for large-scale studies or genomic resources encompassing hundreds of thousands or millions of genomes. Here we introduce gTranslate, a computationally efficient tool that provides improved performance for predicting translation tables for bacteria. We introduce several genomic features that are predictive of bacterial genetic codes and use these to train an ensemble of five machine learning classifiers. Unlike the coding density-based heuristic, which can only distinguish between translation table 11 and a UGA recoding, gTranslate can differentiate between translation tables 11, 4, and 25. During evaluation of gTranslate we identified the first members of the genus *Ca*. Stammera to use genetic code 11 instead of 4 and the first members of the phylum Patescibacteriota to use genetic code 4. These predictions are supported by additional genomic evidence.

## Materials & Methods

### Training and Validation Data

gTranslate was trained and validated using genomes in the Genome Taxonomy Database (GTDB; Parks et al., 2022) with both GTDB and NCBI taxonomic classifications used to establish the expected ground truth for each genome (see *Establishing the Expected Ground Truth* below). The 596,861 genomes in GTDB R09-RS220 were dereplicated to 356,020 genomes by randomly selecting at most 100 genomes per species to mitigate biasing results to species represented by large numbers of genomes. The ground truth was table 11 for 354,299 (99.52%) genomes, table 4 for 1,521 (0.43%) genomes, and table 25 for 200 (0.06%) genomes (**Supp. Table 1**). These genomes had an average completeness of 92.0% (min. = 18.0%) and contamination of 1.3% (max. = 49.1%) as estimated with CheckM2 1.0.2 (Chklovski et al., 2023).

### Independent Test Data

The 116,508 genomes in GTDB R10-RS226 not contained in GTDB R09-RS220 were used as an independent test set to evaluate the expected performance of gTranslate (**Supp. Table 2**), with expected ground truth established as for the training and validation data. The ground truth for these genomes was table 11 for 115,940 (99.51%) genomes, table 4 for 447 (0.38%) genomes, and table 25 for 121 (0.10%) genomes. These genomes had an average completeness of 92.7% (min. = 38.0%) and contamination of 1.3% (max. = 22.9%).

gTranslate was also evaluated on 134,075 genomes from GlobDB release 226 (Speth et al., 2024) after removing genomes that: i) appear in GTDB (143,607 filtered genomes), ii) have an unresolved classification at the rank of order or higher which precludes establishing an accurate ground truth (754 filtered genomes), or iii) are <50% complete, >10% contaminated, or <50 quality (completeness-5×contamination) according to CheckM2 estimates (27,822 filtered genomes) (**Supp. Table 3**). The remaining GlobDB genomes come from studies producing large numbers of genome assemblies that have not been submitted to an International Nucleotide Sequence Database Collaboration (INSDC) repository and thus do not appear in any GTDB release. The expected ground truth was established using the taxonomic classifications provided for these genomes by GlobDB, which are determined using GTDB-Tk (Chaumeil et al., 2022) classifications extended by an automated method for assigning genomes to new taxa. The ground truth for these genomes was table 11 for 133,566 (99.62%) genomes, table 4 for 89 (0.07%) genomes, and table 25 for 420 (0.31%) genomes. These genomes had an average completeness of 85.4% (min. = 50.1%) and contamination of 1.78% (max. = 9.98%).

### Establishing the Expected Ground Truth

The ground truth (i.e. presumed correct) translation table for each genome assembly was established by considering its classification in the GTDB and NCBI taxonomic frameworks (Parks et al., 2025; Schoch et al., 2020). Based on prior information, genomes in the GTDB class JAEDAM01, which contains the two GTDB orders *Absconditabacterales* and BD1-5, were considered to use translation table 25. Note that the GTDB class JAEDAM01 as opposed to class *Gracilibacteria* (Rinke et al., 2013) contains organisms using translation table 25 due to an error in the specification of the type strain, which defines the position of *Gracilibacteria* (*personal communication* with Chris Rinke). Genomes belonging to the GTDB order *Mycoplasmatales* (Razin et al., 1998), the GTDB species *Ca*. Zinderia insecticola (McCutcheon and Moran, 2010), and the *Eggerthellaceae* genera CAVGFB01 or JAUNQF01 (Parks et al., 2025) were considered to use translation table 4. Genomes assigned to the NCBI species *Ca*. Hodgkinia cicadicola (McCutcheon et al., 2009), *Ca*. Nasuia deltocephalinicola (Bennett and Moran, 2013), *Ca*. Stammera capleta (Salem et al., 2017), *Ca*. Organicella extenuate (Williams et al., 2021), and *Ca*. Pinguicoccus supinus (Serra et al., 2020) were also considered to use translation table 4. These species have highly reduced genomes and are not included in GTDB as they lack sufficient phylogenetically informative genes to be robustly placed in the reference trees used by this taxonomic resource. *Fastidiosibacteraceae* strain XS4 (Nakayama et al., 2025), which uses translation table 4, was not included in the analysis as no genome assemblies for the species are currently available in the NCBI Assembly database (Kitts et al., 2016). All other species were considered to use the standard bacterial code (translation table 11).

### Calculation of the Feature Vector

Genomes were transformed into a vector consisting of 8 features that were used by all evaluated machine learning (ML) classifiers:

- *Coding density 11* (CD11): gene coding density when predicting genes with Prodigal (Hyatt et al., 2010) using translation table 11.
- *Coding density 4/25* (CD4): gene coding density when predicting genes with Prodigal using translation table 4 (or equivalently 25), which reassigned the UGA stop coding to either tryptophan (or glycine).
- *Coding density difference*: the difference in coding density between CD4 and CD11 (CD4 – CD11).
- *GC percentage:* percentage of guanine (G) and cytosine (C) nucleotides in a genome.
- *Tryptophan ratio*: log-transformed ratio of UGA to UGG codon counts when predicting genes with Prodigal under translation table 4. The log ratio is clamped between -6 and 5 to remove extreme outliers.
- *Tryptophan magnitude*: log-transformed count of all UGA and UGG tryptophan codons when predicting genes with Prodigal under translation table 4.
- *Glycine ratio:* log-transformed ratio of UGA codon counts to the total glycine codon counts (i.e. codons GGn) when predicting genes with Prodigal under translation table 4. The log ratio is clamped between -10 and 0 to remove extreme outliers.
- *UGG density*: ratio of UGG tryptophan codons to glycine codons (GGn) when predicting genes with Prodigal under translation table 4.

Prodigal was run with default parameters with the exception that i) proteins were not called across ambiguous bases by using the restricted gene prediction (-m), and ii) setting the prediction procedure (-p) to ‘meta’ if a genome assembly consisted of <100 kb. While ML classifiers can learn non-linear decision boundaries that utilize all 8 features, pairwise plots are sufficient to illustrate that these features can discriminate between translation tables (**Supp. Figure 1**). Features were normalized between 0 and 1 using min-max scaling for the k-nearest neighbours and multi-layer perceptron classifiers.

### Evaluated Classifiers and Hyperparameter Value Selection

The translation table for each genome was predicted using seven ML classifiers, the CheckM coding density heuristic (Parks et al., 2015) and Codetta (Shulgina and Eddy, 2021). Specifically, we evaluated AdaBoost, k-nearest neighbours (k-NN), decision tree, random forest, and multi-layer perceptron (MLP) classifiers as implemented in the Scikit-learn 1.6.1 Python package (Pedregosa et al., 2011), and the LightGBM 4.6.0 (Ke et al., 2017) and XGBoost 2.1.4 (Chen and Guestrin, 2016) gradient boosting classifiers. A grid search was performed over the most salient hyperparameters for each of the classifiers (**Supp. Table 4**) with 5-fold cross-validation across 80% of randomly selected GTDB R09-RS220 genomes used to determine the hyperparameter values resulting in the highest mean accuracy across the folds. Performance of the classifiers using the selected hyperparameter values was then validated using 5-fold cross-validation across the full set of GTDB R09-RS220 genomes to evaluate initial classifier performance and to ensure classifiers did not overfit to the 80% of data available during hyperparameter value selection.

Codetta provides evidence for alternative genetic codes and does not directly provide a prediction for a specific translation table. Specifically, alignments of profile hidden Markov models of conserved proteins to codons are used to identify the most likely translation for each codon. We evaluated the accuracy of Codetta to correctly predict translation table 11, 4, or 25 by examining the evidence provided by Codetta for a UGA stop codon reassignment. Specifically, we follow Shulgina and Eddy (2021) and evaluated Codetta assuming either 8 or 34 columns as sufficient support to treat the UGA stop codon as being reassigned to tryptophan or glycine and thus a prediction of translation table 4 or 25, respectively.

The accuracy of the translation table information provided by NCBI, which is strictly based on taxonomic information (taxon ids), was also assessed. Translation table information was obtained from the GenBank Flat File (GBFF) for each genome when available or the translation table associated with an NCBI species classification. NCBI translation table information could only be established for 207,494 (58.3%) of the 356,020 genomes in GTDB R09-RS220 as many genomes lack an NCBI species assignment (**Supp. Table 1**).

### Ensemble of Classifiers

All possible combinations of 3 and 5 ML classifiers were formed into an ensemble with majority voting used to establish the final prediction. Ties, while rare, are possible given that there are 3 possible translation tables. Two types of ties can occur when considering an ensemble of 5 classifiers. The first involves a tie resulting from two table 11 predictions, two table 4 (25) predictions, and a single vote for table 25 (4). In this case, majority vote predicts a reassignment of the UGA stop codon and the tie is resolved by predicting either table 4 or 25. For example, if the 5 votes are 11, 11, 4, 4, and 25 the ensemble classifier will predict table 4. The second type of tie can occur when using an ensemble of 3 or 5 classifiers and occurs when there are an equal number of votes for tables 4 and 25 (e.g. 11, 4, 4, 25, 25). In this case, there is no unambiguous way to resolve the tie between table 4 and 25. Consequently, gTranslate will produce a warning and predict table 4 as this table occurs more frequently in large genome datasets. In practice, this situation is exceedingly rare. It occurred only once across the validation tests with R09-RS220 and did not occur in the final R10-RS226 or GlobDB testing when using the 5-classifier ensemble ultimately selected for use in gTranslate.

### Leave-one-taxon-out Validation

The performance of the 7 ML classifiers was evaluated using a leave-one-taxon-out procedure to validate performance on genomes representing different levels of taxonomic novelty. Specifically, all genomes in a single GTDB taxon were removed from the GTDB R09-RS220 set, classifiers were then trained on the remaining genomes, and prediction results evaluated on the genomes from the removed taxon. This procedure was performed for the ranks of species, genus, and family as removing taxa at broader ranks can result in insufficient training data for translation table 4 and 25.

### Tree Inference

Phylogenetic trees were inferred for lineages with consistently different translation table predictions to the ground truth. A *Ca*. Stammera capleta tree was inferred from a set of 181 core genes identified using Panaroo 1.5.2 (Tonkin-Hill et al., 2020) run in strict mode using a sequence identity threshold of 60%, protein family identity threshold of 50%, length difference threshold of 5%, and core gene threshold of 70%. Genes were aligned using MAFFT 7.505 (Katoh and Standley, 2013) run using the L-INS-i alignment method and --maxiterate set to 1000. A maximum-likelihood tree was inferred with IQ-Tree 2.4.0 (Minh et al., 2020) with the evolutionary model GTR+F+R5 selected using ModelFinder (Kalyaanamoorthy et al., 2017). Support values were determined using 1000 replicates for both the SH approximate likelihood ratio test (SH-aLRT; Anisimova et al., 2011) and ultrafast bootstraps (UFBoot; Hoang et al., 2018). As in previous studies of this species (García-Lozano et al., 2024), the tree was rooted using *Buchnera aphidicola*.

Phylogenetic trees for *Minisyncoccia* genomes in support of the JAKLIH01 family and GCA-2747955 genus having a UGA stop codon reassignment were inferred using the 120 GTDB bacterial marker genes (bac120). These trees were inferred using GlobDB release 226 genomes with families consisting of >10 genomes dereplicated to the 10 highest-quality genomes based on CheckM2 estimates. Bac120 marker genes were identified, aligned and trimmed using the *de novo* inference workflow of GTDB-Tk 2.5.2 with default parameters. Maximum-likelihood trees were inferred with IQ-Tree 2.4.0 with the evolutionary model LG+F+I+R10 selected using ModelFinder for both trees. Support values were determined with 1000 replicates for SH-aLRT and UFBoot. Trees were rooted to conform to the GTDB R10-RS226 reference tree topology.

### Calculation of Genome Properties

Genomes were annotated with Prokka 1.14.6 (Seemann, 2014) using its bacterial reference databases. Aragorn 1.2.38 (Laslett, 2004) was used internally by Prokka to identify tRNA genes. Prokka annotations were inspected to identify tRNA^Trp^(UCA), tRNA^Gly^(UCA) and peptide chain release factor 2 (RF2, *prfB*) genes. %GC content and genome size were determined using CheckM2 1.0.2.

## Results

### Selection of Classifier Hyperparameter values

A grid search was conducted over the most salient hyperparameters for seven ML classifiers under consideration for inclusion in gTranslate, with performance assessed using 5-fold cross-validation over 284,816 (80%) of the 356,020 GTDB R09-RS220 genomes (**Supp. Table 4**). Classifier performance varied substantially across parameters highlighting the importance of conducting hyperparameter value selection (**Figure 1**). However, all classifiers with the exception of MLP had similar performance over a range of hyperparameter values indicating robustness to value selection. The combination of hyperparameter values resulting in the highest accuracy (fewest misclassifications) cumulatively across the 5 folds was selected for each classifier.

**Figure 1.**
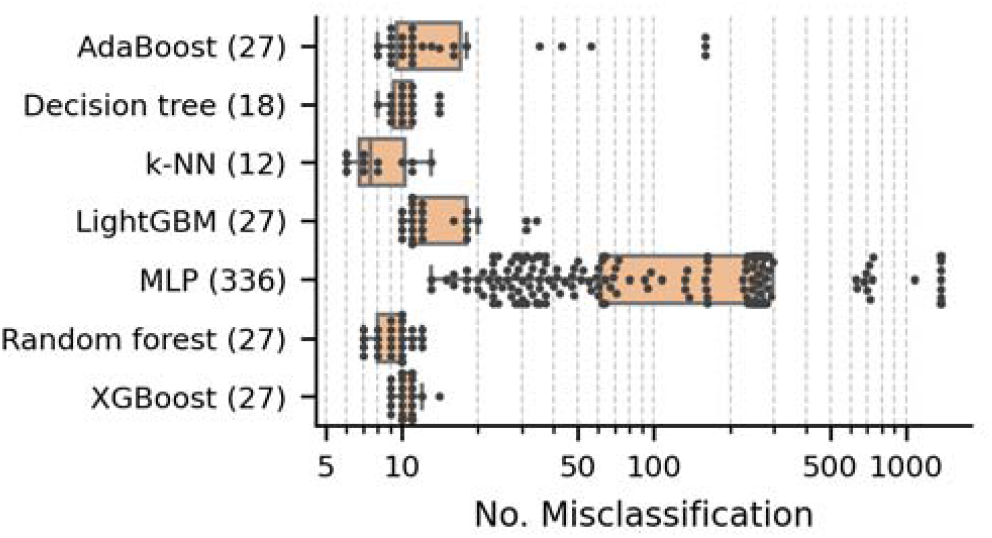
Distribution of misclassifications (log scale) on GTDB R09-RS220 genomes across combinations of hyperparameter values for seven machine learning classifiers. The total number of parameter combinations tested is indicated in parentheses. The box plots represent the median and interquartile range of performance, while the overlaid swarm plots depict the results for individual combinations.

### Classifier Performance using Cross Validation

We evaluated the performance of the seven ML classifiers and 56 ensembles of these classifiers using 5-fold cross validation across the 356,020 genomes in GTDB R09-RS220. This includes the 20% of genomes withheld during hyperparameter selection to ensure that the classifiers are generalizing to new genomes. Performance varied across the classifiers with k-NN achieving the best results with only 15 misclassifications and the decision tree performing the worst with 30 misclassifications (**Figure 2A; Supp. Figure 2**). The best ensemble classifiers were able to reduce the number of misclassifications to 14, with many different combinations of classifiers outperforming all individual classifiers except k-NN (**Supp. Table 5**). All classifiers and classifier ensembles had better performance than the CheckM coding density heuristic (101 misclassifications) or Codetta requiring either 8 (444 misclassifications) or 34 (78 misclassification) columns supporting a UGA stop codon reassignment (see *Materials & Methods*; **Figure 2A; Supp. Table 1**). NCBI translation table data was available for 207,495 (58.3%) of the genomes, and 62 of these were found to disagree with the expected ground truth (**Supp. Table 1**). This includes: i) 28 genomes classified by NCBI as *Candidatus* Gracilibacteria with assembly data indicating table 25, but GTDB assignments to taxa that use table 11, ii) 16 genomes classified by NCBI only as Bacteria were assigned to table 11, whereas GTDB assigns these taxa to table 4 (*Mycoplasmatales*) or table 25 (JAEDAM01), and iii) 6 genomes classified by NCBI to the family *Mycoplasmoidaceae* were assigned to translation table 11, which should be table 4 based on the taxon. Notably, both the CheckM heuristic and Codetta results with 34 columns agree with the ground truth for all but two of these genomes indicating that the translation table data provided by NCBI is erroneous in most of these cases.

**Figure 2.**
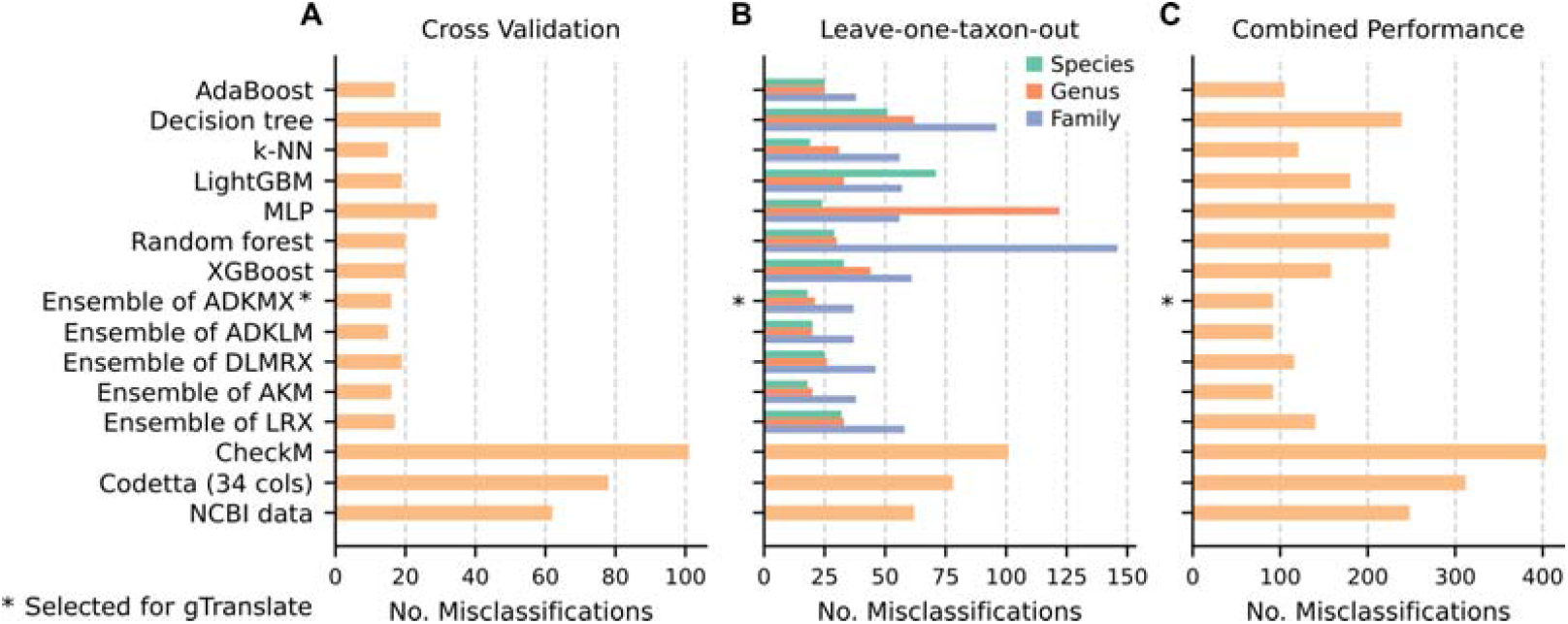
Validation performance on GTDB R09-RS220 genomes. The number of misclassified genomes is provided for both (**A**) cross validation and (**B**) leave-one-taxon-out testing methodologies (evaluated at 3 taxonomic ranks for the ML classifiers), and (**C**) the combined performance over these two test methodologies. Results are given for individual ML classifiers, the ensemble of classifiers resulting in the best and worst combined performance, the CheckM coding density heuristic, Codetta, and the translation table data provided at NCBI. Codetta results are shown for 34 required columns of support as results requiring only 8 columns were substantially worse (444 misclassifications). Note that the single leave-one-taxon-out misclassifications for the non-ML methods were counted 3 times to calculate the combined performance to be comparable to the ML taxon-specific results. Results for all ensemble classifiers are provided for the cross-validation test in Supp. Figure 2 and leave-one-taxon-out test in Supp. Figure 3. For the ensemble classifiers, A = AdaBoost, D = decision tree, K = k-NN, L = LightGBM, M = MLP (multi-layer perceptron), R = random forest, and X = XGBoost.

### Classifier Performance on Taxonomically Novel Genomes

The ML classifiers and classifier ensembles were validated using a leave-one-taxon-out procedure to evaluate performance on genomes representing different levels of taxonomic novelty. This was performed by removing a single GTDB taxon from the GTDB R09-RS220 validation set, training on the remaining genomes, and then evaluating the predictions on genomes from the removed taxon. Performance of individual classifiers and ensembles generally decreases with taxonomic novelty (**Figure 2B; Supp. Figure 2**), and there was insufficient training data to evaluate classifier performance above the rank of family. For example, leaving out the 1,467 genomes from the order *Mycoplasmatales* leaves only 52 genomes that use translation table 4. Performance of individual classifiers varied substantially; for example, at the rank of species k-NN resulted in only 19 misclassifications compared to LightGBM at 71 misclassifications (**Supp. Table 5**). Notably, ensembles of classifiers produced more consistent results. At the species level, the best 5 ensemble classifier had 18 misclassifications, while the worst had 26. Since the CheckM heuristic, Codetta, and translation table data at NCBI are not dependent on a training set, their predictions are consistent across this validation procedure and produce the same misclassifications as reported in *Performance using Cross Validation*. Notably, all ensemble classifiers outperform these methods.

### Selecting a Classifier Ensemble

The combined results from the cross validation (**Figure 2A**) and leave-one-taxon-out (**Figure 2B**) were used to select the ML classifier(s) for gTranslate. Ensembles of 3 or 5 classifiers consistently outperformed individual classifiers (**Figure 2C; Supp. Figures 2** and **3**). The best performing individual classifiers were AdaBoost with 105 combined misclassifications and k-NN with 121 misclassifications (**Supp. Table 5**). By contrast, there were three classifier ensembles that produced only 92 misclassifications: i) AdaBoost, k-NN, and MLP and ii) AdaBoost, Decision tree, k-NN, MLP, with either XGBoost or LightGBM. We selected the ensemble of 5 classifiers with XGBoost instead of the equally performing 3-classifier ensemble as it provides gTranslate users with additional information since results are provided for each ML classifier. Preference was given to XGBoost over LightGBM as this 5-classifier ensemble produced slightly superior performance on the leave-one-taxon-out results at the rank of species, a situation which we expect to occur frequently in real data.

### Performance of gTranslate on GTDB Genomes

The expected (test) performance of gTranslate was evaluated by training on the dereplicated set of 356,020 GTDB R09-RS220 genomes and assessing classification accuracy on 116,508 genomes introduced in GTDB R10-RS226. These test genomes represent all new genomes between two annual releases of the GTDB, many of which are from new taxa (**Table 2; Supp. Table 6**). This test set is expected to be indicative of the taxonomic novelty of genomes that gTranslate will be applied to, and hence real-world performance, as gTranslate will be retrained annually with each GTDB release.

**Table 2.**
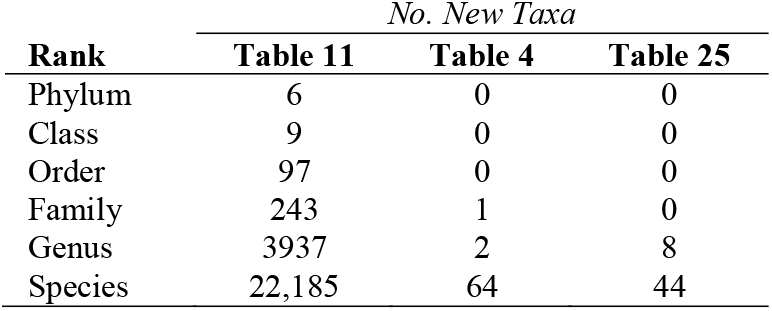
Taxonomic Novelty of the 116,508 GTDB R10-RS226 Test Genomes.

gTranslate correctly predicted the translation table of 116,502 (99.995%) of the 116,508 test genomes with only 6 differing from the expected ground truth (**Supp. Table 2**). Of these, 5 are *Ca*. Stammera capleta genomes that are expected to use table 4 but predicted to use table 11 by gTranslate. Prediction of table 11 is strongly supported by all classifiers comprising the gTranslate ensemble as well as the CheckM heuristic and Codetta. Notably, Codetta strongly supports a UGA stop to tryptophan reassignment for all *Ca*. S. capleta genomes except for these 5. Furthermore, all 5 genomes retain *prfB* (peptide chain release factor RF2), which is typically lost in genomes using genetic code 4 (Bezerra et al., 2015), including in the majority of *Ca*. S. capleta genomes (**Figure 3**). Unlike most species using genetic code 4, *Ca*. S. capleta genomes do not have a tRNA^Trp^(UCA) codon for recognizing UGA as tryptophan (Manzano-Marín et al., 2023), which precludes using this gene as further genomic evidence in support of *Ca*. S. capleta genomes using genetic code 4 or 11. Nonetheless, most genomic evidence supports the 5 genomes using genetic code 11 as predicted by gTranslate. Moreover, these genomes form a monophyletic clade basal to all other *Ca*. S. capleta (**Figure 3**), a placement observed in previous studies of this species (García-Lozano et al., 2024). This suggests that codon reassignment may have occurred after the basal clade diverged, noting however that genome GCA_036689315.1, which uses genetic code 4, is superficially basal to this group (i.e. without support; **Figure 3**).

**Figure 3.**
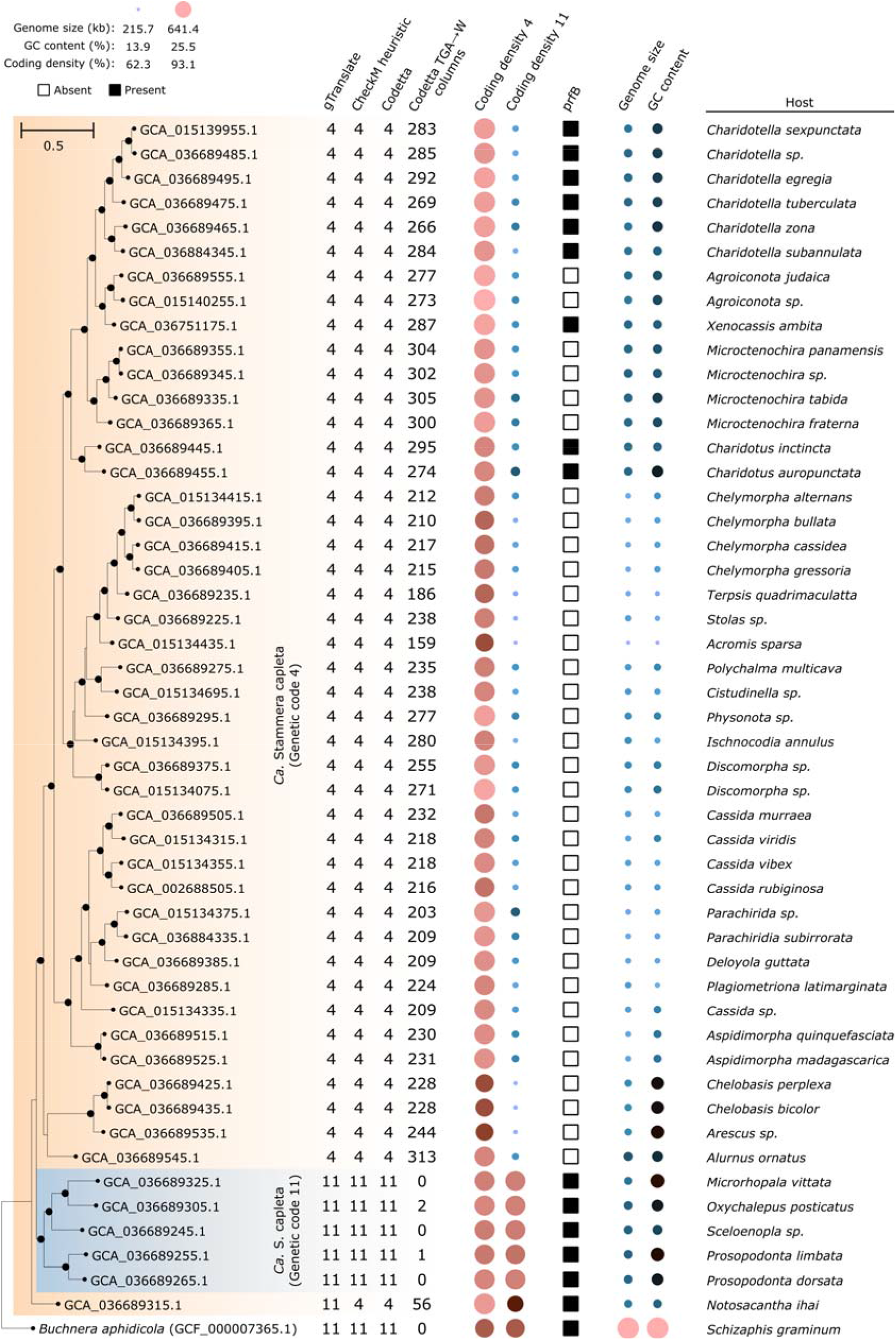
Maximum-likelihood phylogenetic tree inferred from alignment of core *Ca*. Stammera capleta genes with IQ-Tree and the GTR+F+R5 evolutionary model. The 5 *S. capleta* genomes with strong evidence of using genetic code 11 are shown in blue shading. Black dots indicate well supported internal nodes - UFBoot ≥95% and SH-aLRT ≥80%. *prfB* = peptide chain release factor RF2.

The other discrepant gTranslate prediction is for the *Patescibacteriota* genome GCA_034717395.1, which belongs to the GTDB species JAYELJ01 sp034717395 (**Supp. Table 2**). This species is expected to use table 11 but predicted to use table 25. Again, this prediction is strongly supported with 4 of the 5 gTranslate classifiers making this prediction, the CheckM heuristic also supporting a reassignment of the UGA stop codon, and Codetta identifying 152 columns with UGA aligned to a tryptophan.GCA_034717395.1 is the only genome in the GTDB order JAYELJ01, and its position on a long branch with minimal support suggests the possibility that the genome is incorrectly placed within the GTDB reference tree and should be assigned to a *Patescibacteriota* order expected to use translation table 25 (**Supp. Figure 4**).

These results suggest that gTranslate may be making correct predictions for all 116,508 test genomes. The CheckM heuristic is in agreement with gTranslate for the 6 contested cases above, however makes an additional 8 incorrect predictions (**Figure 4A**). All 8 of these misclassifications appear to be legitimate errors and, unlike gTranslate, CheckM is unable to distinguish between genomes using table 4 or 25 (**Supp. Table 2**). Of these misclassifications, 7 are incorrectly predicting a UGA stop codon reassignment. The other misclassification is predicting the use of table 11 for a *Mycoplasmatales* genome. Codetta makes 111 or 6 additional misclassifications when requiring 8 or 34 columns of support for a UGA reassignment, respectively. All misclassifications are erroneous UGA reassignments. The 6 misclassified genomes with ≥34 columns of support are distinct from the misclassifications made by the other methods (**Figure 4A**), are predicted in 6 separate phyla, and include genomes from species that are unlikely to have an unrecognized UGA stop codon reassignment such as *Salmonella enterica* and *Lactococcus cremoris*.

**Figure 4.**
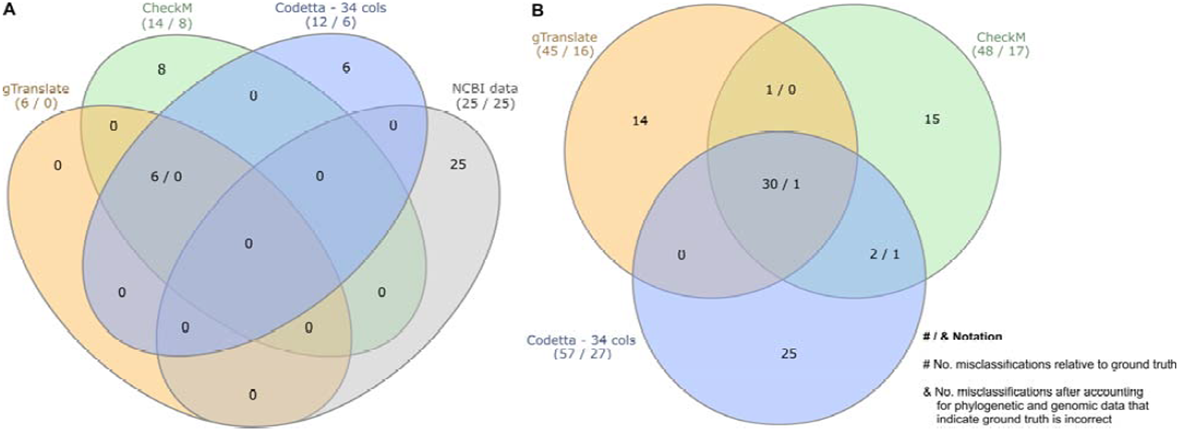
Venn diagram showing overlap in misclassifications between gTranslate, CheckM coding density heuristic, Codetta requiring 34 columns of support, and NCBI translation table data on the **(A)** 116,508 GTDB R10-RS226 and **(B)** 134,075 GlobDB release 226 test genomes. Numbers in parentheses indicate total number of putative misclassifications relative to ground truth and after considering additional evidence suggesting errors in the ground truth (#/& notation where these numbers differ). Diagram generated with InteractiVenn (Heberle et al., 2015).

NCBI provides translation table information for 83,262 (71.5%) of the 116,508 test genomes (**Supp. Table 2**). This information differs from the ground truth for 25 genomes, 24 of which are assigned to the phylum *Candidatus* Gracilibacteria by NCBI (class *Gracilibacteria* in GTDB R09-RS220). The remaining genome (GCA_035437565.1) is only classified as Bacteria at NCBI, and as such is predicted to use translation table 11. However, this genome belongs to the class JAEDAM01 in GTDB R09-RS220, a taxon comprised of genomes using translation table 25. Notably, gTranslate, the CheckM heuristic, and Codetta agree with the ground truth for all 25 of these genomes (**Figure 4A**).

### Performance of gTranslate on GlobDB Genomes

Test performance of gTranslate was further evaluated by assessing classification accuracy on 134,075 genomes from GlobDB release 226. The GlobDB test genomes are derived from metagenomic studies not submitted to INSDC, thus representing additional taxonomic novelty relative to GTDB R10-RS226 (**Table 3; Supp. Table 7**). GlobDB is updated with each GTDB release and represents a common use case for gTranslate where large number of genomes of varying taxonomic novelty must be processed.

**Table 3.** Taxonomic Novelty of the 134,075 GlobDB Test Genomes.

| Rank | No. New Taxa |  |  |
| --- | --- | --- | --- |
|  | Table 11 | Table 4 | Table 25 |
| Phylum | 24 | 0 | 0 |
| Class | 119 | 0 | 0 |
| Order | 530 | 0 | 0 |
| Family | 3,643 | 10 | 13 |
| Genus | 25,776 | 26 | 99 |
| Species | 90,619 | 32 | 228 |

gTranslate correctly predicts the translation table of 134,030 (99.97%) of the GlobDB test genomes (**Supp. Table 3**). However, like the results on the GTDB R10-RS226 test genomes, the majority of the 45 putative misclassifications appear to be errors in the ground truth due to previously unrecognized stop code reassignments or taxonomic misclassifications (**Figure 4B**). The most notable of these are 12 genomes from the JAKLIH01 family and 10 genomes from the GCA-2747955 genus within the class *Minisyncoccia* where genomic evidence indicates a reassignment of the UGA stop codon to tryptophan (**Figure 5; Supp. Table 8**). These 22 genomes have large numbers of columns where UGA aligns with tryptophan in conserved proteins, mostly have an identified tRNA^Trp^(UCA) gene (i.e. the anticodon of UGA), and lack a *prfB* gene consistent with organisms using a reassigned UGA stop codon. These are the first organisms in the *Patescibacteriota* with a predicted UGA-to-tryptophan reassignment making this the first bacterial phylum to use three translation tables.

**Figure 5.**
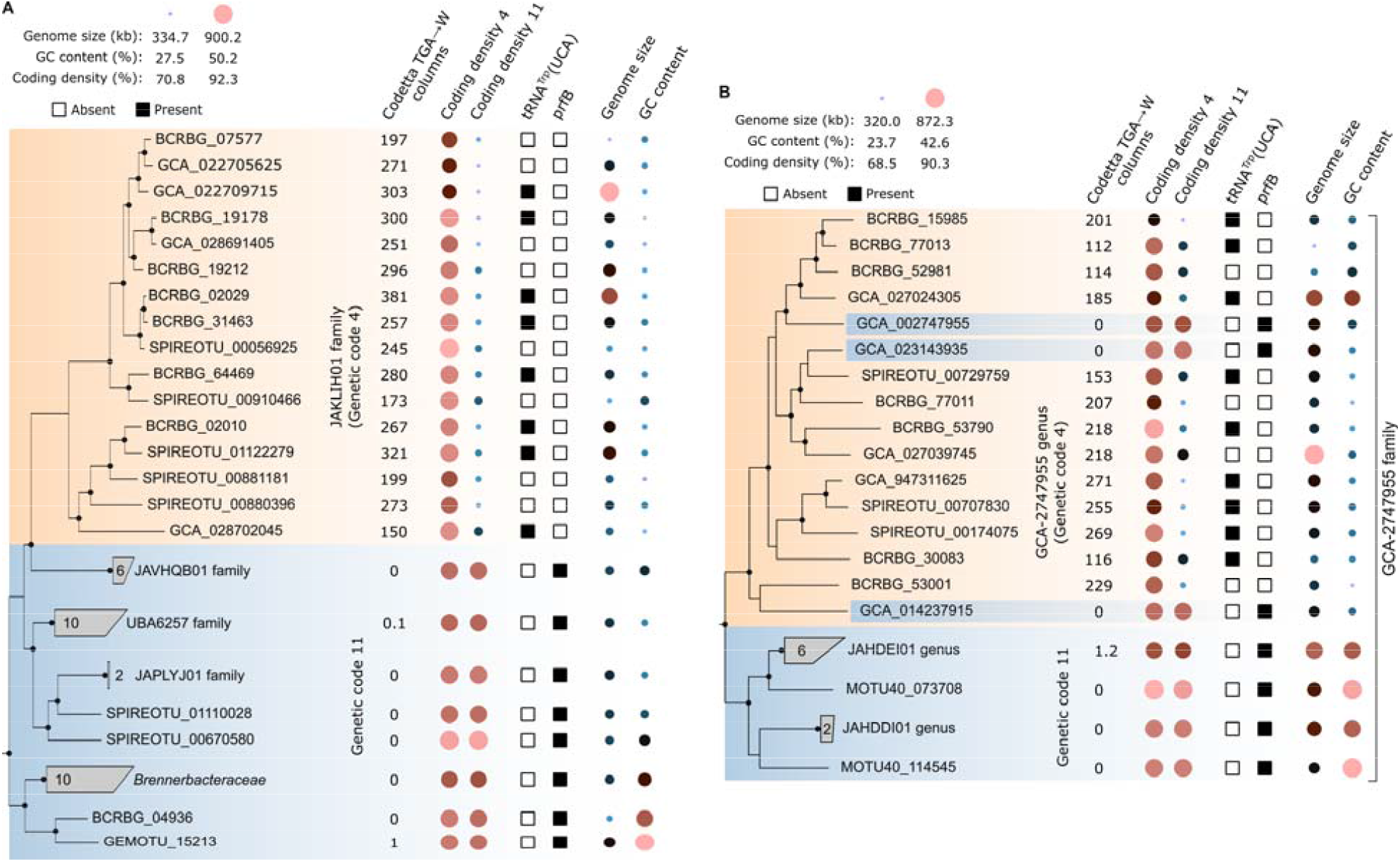
Genomic evidence supporting the use of translation table 4 in the class *Minisyncoccia* belonging to the bacterial phylum *Patescibacteriota*, specifically (**A**) the family JAKLIH01 and (**B**) genus GCA-2747955. The maximum-likelihood trees were inferred from a set of 120 bacterial marker genes using IQ-Tree with the LG+F+I+R10 evolutionary model. Data is provided for the 16 JAKLIH01 and 14 GCA-2747955 genomes in GlobDB release 226 along with genomes from closely related families (**A**) or genera (**B**). The 3 genomes in the GCA-2747955 genus shown with blue highlighting still use genetic code 11 despite being in a lineage otherwise comprised of genomes with the UGA stop codon reassigned to tryptophan. Genomes with identifiers that do not start with GCA are part of the GlobDB test set, whereas genomes starting with GCA are part of GTDB. Taxa were dereplicated to at most 10 genomes, with taxa represented by a single genome labelled using the GlobDB genome identifier. Genomic properties for taxa with multiple genomes indicate the average across these genomes. The *prfB* gene (peptide chain release factor RF2) was identified in all genomes using genetic code 11 except for a single *Brennerbacteraceae* genome. Black dots on internal branches indicate strong support with UFBoot ≥95% and SH-aLRT ≥80%.

There are also two genomes belonging to the *Eggerthellaceae* with a ground truth of table 11 predicted to use table 4 by gTranslate, the CheckM heuristic, and Codetta (**Supp. Table 3**), consistent with our recent report of UGA-to-tryptophan reassignment in members of this bacterial family (Parks et al., 2025). Similar to the GCA_034717395.1 genome in the GTDB R10-RS226 test set, there are 5 genomes assigned to the *Patescibacteriota* order GCA-2401425 with a ground truth of 11, but genomic evidence that they use table 25 (**Supp. Table 3**) and thus are likely taxonomically misclassified (**Supp. Figure 4**). Finally, there is a single *Burkholderiaceae* genome that was assigned to a novel genus by GlobDB and thus has a default ground truth of table 11. However, this genome is assigned to the genus *Zinderia* when using GTDB-Tk with the GTDB R10-RS226 reference data, contains a tRNA^Trp^(UCA) gene, lacks a *prfB* gene, is classified to table 4 by the CheckM heuristic and has 30 columns identified by Codetta in support of a UGA-to-tryptophan reassignment (**Supp. Table 3**). This suggests that this genome should be classified as *Z. insecticola* and that it uses translation table 4.

gTranslate predictions appear to be legitimately incorrect for 13 genomes in the *Patescibacteriota* class JAEDAM01 (**Supp. Table 3**). These genomes are all expected to use table 25 based on their taxonomic classification, and the Codetta results support this assignment. However, gTranslate predicts table 4 for 11 of these genomes and table 11 for the remaining 2 genomes. In addition, there are two *Mycoplasmatales* genomes that gTranslate incorrectly predicts as using table 25 or 11 instead of table 4 (**Supp. Table 3**). Surprisingly, the CheckM heuristic and Codetta both predict table 11 along with gTranslate for the BCRBG_74280 *Mycoplasmatales* genome; however, this genome has a tRNA^Trp^(UCA) gene and lacks a *prfB* gene suggesting the ground truth is correct. There are also two *Minisyncoccia* genomes that are predicted to have a reassigned stop codon according to the CheckM heuristic and Codetta (**Supp. Table 3**). Genomic evidence supports a stop reassignment for one of these genomes and indicates it uses table 4 as identified for other *Minisyncoccia* lineages in this study. The other reassignment is to table 25 but the absence of a tRNA-Gly(UCA) and presence of the *prfB* gene do not support this recoding, making it more likely that this misclassification is due to the relatively poor quality of the genome assembly (74.4% complete; 4.2% contamination).

The 15 and 25 misclassifications made only by the CheckM heuristic or Codetta, respectively, appear to be legitimate errors (**Supp. Table 3**). All 15 of the misclassifications exclusive to the CheckM heuristic are for genomes with a predicted UGA stop codon reassignment. Notably, these genomes have abnormally low coding density when genes are called using table 4/25 (mean = 81.3%) or table 11 (75.6%), that may be in part due to a high number of contigs in the assemblies (mean = 506). Most of these genomes have a *prfB* gene (11 of 15) and all lack a tRNA-Trp(UCA) or tRNA-Gly(UCA) gene indicating that the ground truth is correct. The 25 misclassifications exclusive to Codetta also all have a ground truth of table 11 with 23 and 2 genomes predicted to use tables 25 and 4, respectively. Most of these genomes (21 of 25) have <100 columns supporting a stop codon reassignment, mostly have a *prfB* gene (22 of 25), and all genomes lack a tRNA-Trp(UCA) or tRNA-Gly(UCA) gene indicating that the ground truth of table 11 is likely correct.

## Discussion

gTranslate provides robust predictions of translation tables used by bacterial genomes, making only 16 misclassifications (99.994% accuracy) across the 250,583 test genomes in GTDB R10-RS226 (Parks et al., 2025) and GlobDB release 226 (Speth et al., 2024). Most misclassifications (12 of 16) were a result of incorrectly distinguishing between tables 4 and 25 that both reassign the UGA stop codon, resulting in an accuracy of 98.9% across the 1,077 genomes that do not use the standard bacterial translation table 11 (**Supp. Tables 2** and **3**). Notably, gTranslate has better accuracy than the taxonomically-defined ground truth as five *Ca*. Stammera capleta (**Figure 3**) and 22 *Minisyncoccia* genomes (**Figure 5**) have genomic evidence indicating the use of a translation table that conflicted with taxonomic expectations. In addition, we attributed nine putative gTranslate misclassifications to inaccurate taxonomic classifications either due to lack of taxonomic resolution provided by automated methods (**Supp. Table 3**) or misplacement in reference phylogenies (**Supp. Figure 4**). The challenge in using taxonomic classifications to establish the translation table for genomes is further illustrated by information provided at NCBI being incorrect for 25 of the 83,262 (99.97% accuracy) GTDB R10-RS226 genomes where NCBI translation table information was available.

gTranslate outperforms the CheckM coding density heuristic and Codetta that had 25 and 33 misclassifications across the 250,583 test genomes, respectively. Furthermore, gTranslate can distinguish between tables 4 and 25, whereas the CheckM heuristic can only indicate that a UGA stop codon reassignment has occurred. This is particularly notable as only 4 of the 16 misclassifications made by gTranslate are due to incorrectly distinguishing table 11 from a UGA stop codon reassignment. While Codetta is treated as a classifier for predicting translation tables 4, 11, and 25 in this manuscript (see *Methods*), it is a more general tool that aims to identify the most likely amino acid encoded by each codon (Shulgina and Eddy, 2021). The Codetta results reported here only consider reassignment of the UGA stop codon and do not consider any additional insights (or misclassifications) that may arise by considering the reassignment of other codons. gTranslate is a focused tool that allows for more computationally efficient prediction of established translation tables with higher accuracy than Codetta. For reference, gTranslate is ∼100x times more computationally efficient than Codetta requiring only ∼20 seconds per genome per CPU on an Intel Xeon E5-2640 v4 2.40 GHz (released in 2016). In practical terms, Codetta is challenging to apply to large datasets as illustrated by it requiring 5 weeks of computation when using 96 E5-2640 CPUs to process the 116,508 new genomes in GTDB R10-RS226.

The results on the GTDB and GlobDB test genomes indicate gTranslate can accurately predict the translation table of genomes with varying levels of taxonomic novelty (**Tables 2** and **3**). Nonetheless, the performance of gTranslate is best when it is trained on genomes similar to those for which it is making predictions as illustrated by the leave-one-taxon-out validation analysis (**Figure 2B; Supp. Figure 2**). A benefit of basing gTranslate on GTDB genomes is that this resource will be updated annually allowing for regular retraining of gTranslate with each GTDB release. Annual updating will also allow incorporation of new stop codon reassignments such as those reported in this and other recent studies (Nakayama et al., 2025; Parks et al., 2025).

Direct genomic evidence in support of a UGA stop codon reassignment such as the presence of a tRNA^Trp^(UCA) (McCutcheon et al., 2009) or tRNA^Gly^(UCA) (Campbell et al., 2013), or lack of *prfB* (Bezerra et al., 2015) is considered throughout this study. While these are key molecular signatures of a reassigned UGA stop codon, they are not well suited for predicting the translation table used by individual genomes. This is exemplified by strains of *Ca*. Stammera capleta (Salem et al., 2017) that use genetic code 4 despite always lacking a tRNA^Trp^(UCA) and containing a *prfB* in some assemblies (**Figure 3**). More generally, translation table prediction must be robust to partial genome assemblies as these predictions are an initial step in bioinformatic programs such as CheckM (Chklovski et al., 2023) and GTDB-Tk (Chaumeil et al., 2022), which are applied to large numbers of genomes, including large-scale studies consisting of incomplete metagenome-assembled genomes. Molecular signatures are therefore best interpreted in a phylogenomic context, where evidence can be evaluated across multiple (potentially incomplete) genomes alongside additional genomic indicators, such as UGA codon alignment in conserved proteins and coding density under different translation tables.

gTranslate is currently limited to prokaryotic genomes and predicting translation tables 11, 4, or 25 where UGA is a stop codon or translated as tryptophan or glycine, respectively. However, other reassignments have been computationally inferred and experimentally verified in some cases. Application of Codetta to over 250,000 prokaryotic genomes revealed a number of putative arginine sense codon reassignments (Shulgina and Eddy, 2021), though these have not yet been experimentally verified (Zürcher et al., 2022). More recently, it has been computationally predicted that a phylogenetically sporadic set of archaeal genomes likely have genome-wide reassignment of the UAG stop codon to pyrrolysine, which has been confirmed by proteomic analysis for two genomes (Kivenson et al., 2025). Notably, this UAG-to-pyrrolysine reassignment is more difficult to detect as it does not result in reduced coding density when genes are identified using translation table 11 as observed for UGA stop codon reassignments. Methods have also been proposed for handling specific fungal (Mühlhausen and Kollmar, 2014) or mitochondrial reassignments (Abascal et al., 2006; Noutahi et al., 2017). These all represent promising directions for future development of gTranslate that would build upon the approach developed in this study. For example, we anticipate incorporating identification of genome-wide UAG stop codon to pyrrolysine reassignments in the next release of gTranslate.

## Conclusion

gTranslate provides a machine learning-based approach for the accurate prediction of prokaryotic translation tables. It improves upon the coding density-based heuristic used by existing bioinformatics software by being able to distinguish between translation tables 4 and 25. By utilizing an ensemble of five classifiers, gTranslate achieves >99.99% accuracy across diverse test datasets while having approximately 100 times greater computational efficiency than Codetta. Use of gTranslate led to the discovery of a basal *Ca*. Stammera capleta lineage that uses the standard bacterial genetic code and the first instances of UGA-to-tryptophan reassignment in the *Patescibacteriota* class *Minisyncoccia*. These findings identify *Patescibacteriota* as the first bacterial phylum to use tables 4, 11, and 25. Although gTranslate is currently limited to the most prevalent prokaryotic codes, we anticipate that annual retraining with GTDB updates will facilitate the future incorporation of novel variations, such as the recent genome wide UAG-to-pyrrolysine reassignment identified in Archaea. Ultimately, gTranslate offers a scalable and robust solution for the high-throughput prediction of translation tables for prokaryotic genomes, ensuring that genetic code variation is accurately captured in large-scale studies and genomic repositories.

## Data Availability

gTranslate is implemented in Python and licensed under the GNU General Public License v3.0. Source code and tool documentation are available at https://github.com/cmc-aau/gTranslate.

## Acknowledgements

We thank Brian Kemish for systems engineering support. We also gratefully acknowledge the researchers who expended significant effort in collecting, sequencing, and depositing the genome assemblies analyzed in this study. Claude Opus 4.6 was used to help generate the Python scripts that produced Figure 1, Figure 2, and Supplementary Figure 1.

## information

This study was funded by research grants from Novo Nordisk Foundation (NNF25SA0105111 to P.H.), Australian Research Council Discovery Project (DP220100900 to P.H.), and strategic funding from The University of Queensland.

