## Supplementary Figures for "gTranslate: rapid and accurate translation table prediction for prokaryotic genomes"

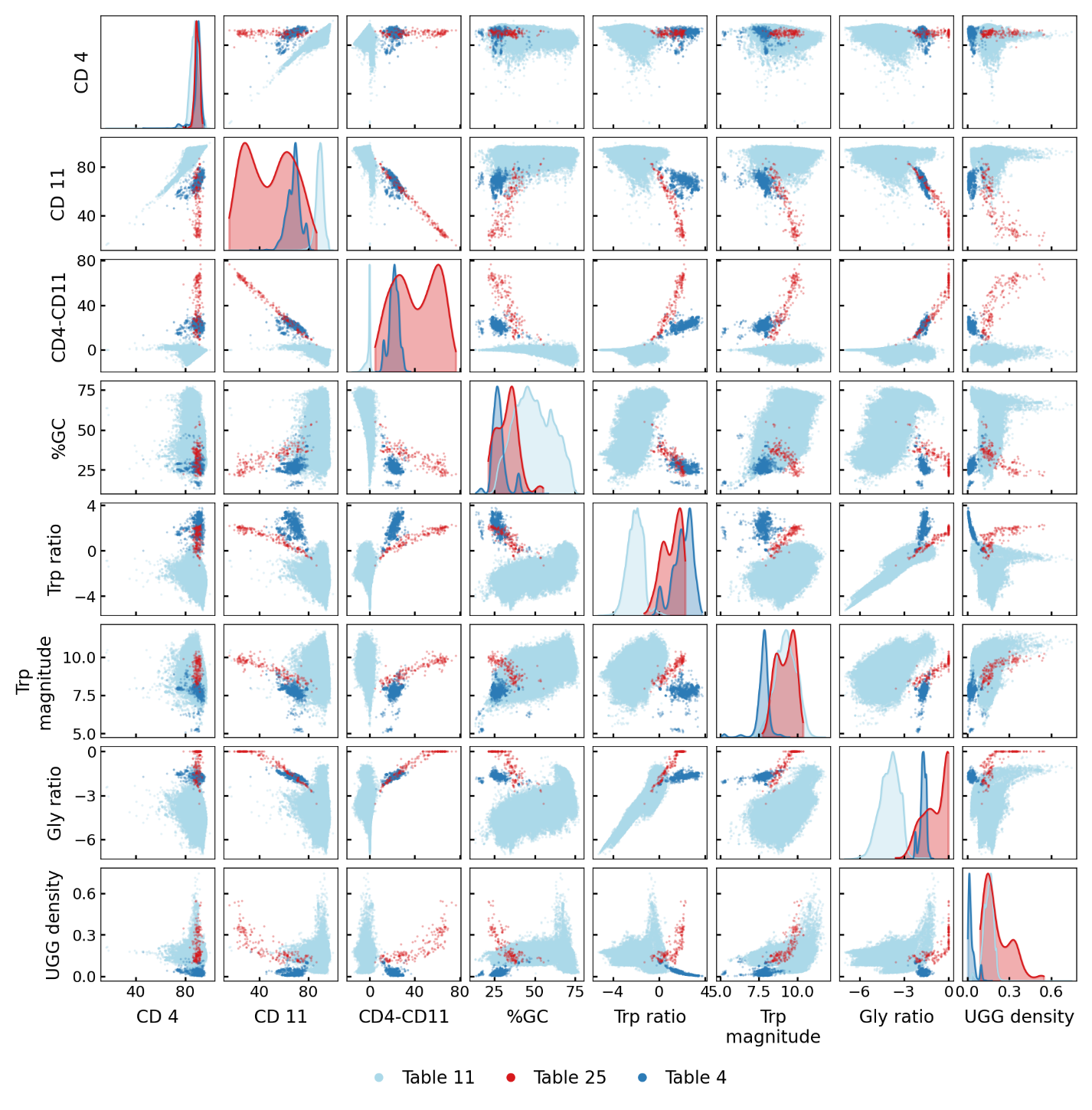

**Supp. Figure 1**. Pairwise scatter plot of 8 genomic features computed for 356,018 prokaryotic genomes from GTDB R09-RS220. Off-diagonal panels display scatter plots for each pair of features, with points coloured by their assigned translation table: table 11 (light blue), table 25 (red), and table 4 (dark blue). Diagonal panels display kernel density estimates (KDEs) for each feature, stratified by translation table and independently normalised to a peak density of one to enable visual comparison across groups despite substantial differences in group size. Feature abbreviations: CD 4 = coding density under translation table 4 (equivalently 25); CD 11 = coding density under translation table 11; CD4-CD11 = difference in coding density between tables 4 and 11; Trp= tryptophan; Gly = glycine.

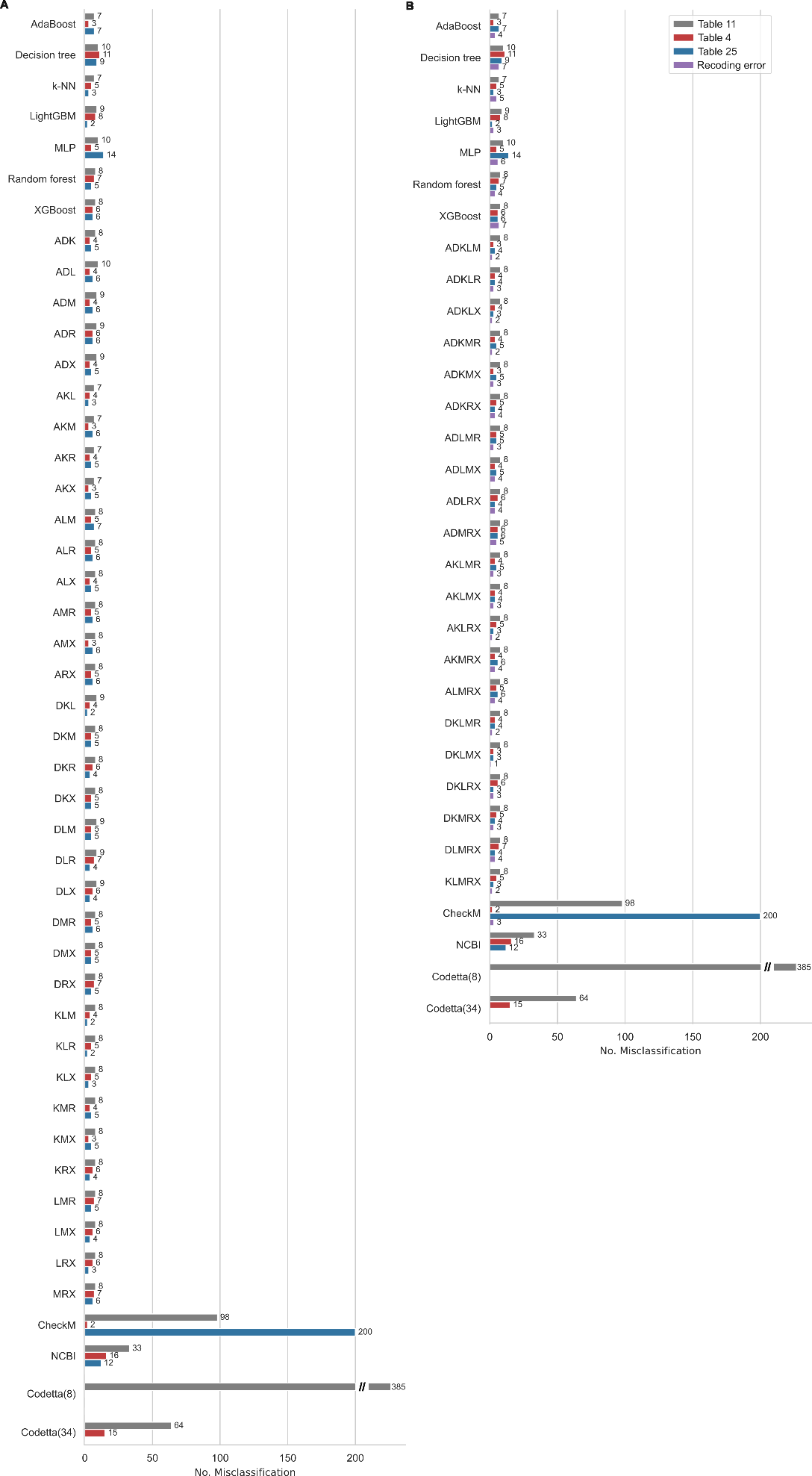

**Supp. Figure 2**. Cross validation performance of the 7 ML classifiers, ensembles of 3 (**A**) and 5 (**B**) classifiers, CheckM heuristic, Codetta with 8 and 34 columns of support, and NCBI. Results indicate the number of misclassifications across all 5 folds made on genomes with a ground truth of table 11 (grey), table 4 (red), or table 25 (blue). Recoding misclassifications (purple; **B**) indicate cases where the ground truth is table 4 or 25 but the classifier predicts table 11.

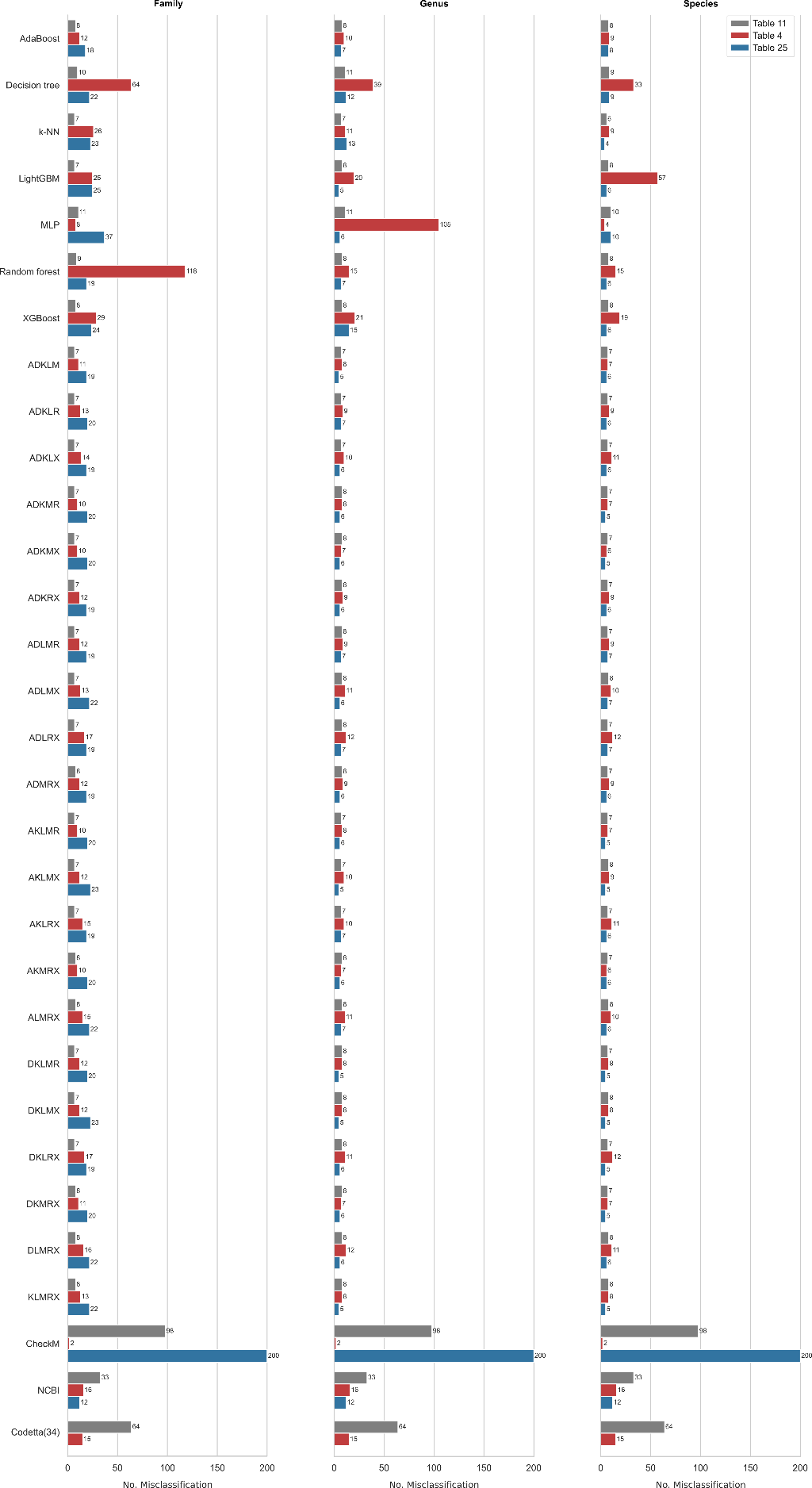

**Supp. Figure 3**. Leave-one-taxon-out performance at the rank of family, genus, and species across 7 ML classifiers, ensembles of 5 classifiers, CheckM heuristic, Codetta with 34 columns of support, and NCBI. Results indicate the number of misclassifications across on genomes with a ground truth of table 11 (grey), table 4 (red), or table 25 (blue).

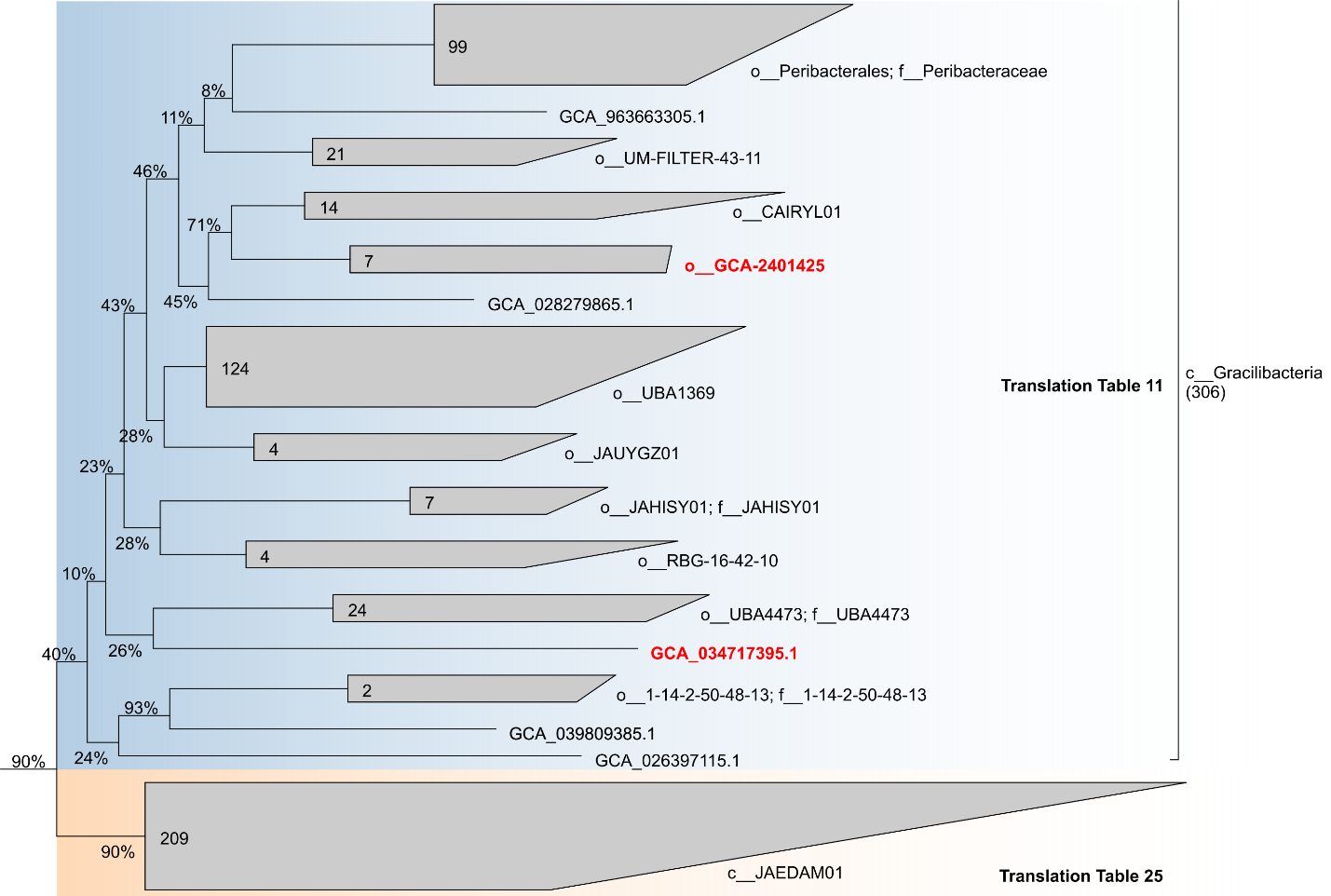

**Supp. Figure 4**. GTDB R10-RS226 reference bacterial tree illustrating the position of the JAYELJ01 sp034717395 genome GCA_034717395.1 and genomes in the order GCA-2401425 (shown in red). The ground truth for these genomes is translation table 11 because of their position within the class Gracilibacteria. gTranslate, CheckM, and Codetta all predict these genomes use translation table 25 as would be expected for a genome in the class JAEDAM01, which is sister to the Gracilibacteria. In addition, all these genomes lack the translational release factor RF2 gene. The relatively low non-parametric bootstrap support values between these genomes and JAEDAM01 suggests they may be incorrectly placed within the GTDB reference tree and could belong to class JAEDAM01.
